# TMEM106B C-terminal fragments aggregate and drive neurodegenerative proteinopathy

**DOI:** 10.1101/2024.06.11.598478

**Authors:** Ruben Riordan, Aleen Saxton, Pamela J. McMillan, Rebecca L Kow, Nicole F. Liachko, Brian C. Kraemer

## Abstract

Genetic variation in the lysosomal and transmembrane protein 106B (TMEM106B) modifies risk for a diverse range of neurodegenerative disorders, especially frontotemporal lobar degeneration (FTLD) with progranulin (PGRN) haplo-insufficiency, although the molecular mechanisms involved are not yet understood. Through advances in cryo-electron microscopy (cryo-EM), homotypic aggregates of the C-Terminal domain of TMEM106B (TMEM CT) were discovered as a previously unidentified cytosolic proteinopathy in the brains of FTLD, Alzheimer’s disease, progressive supranuclear palsy (PSP), and dementia with Lewy bodies (DLB) patients. While it remains unknown what role TMEM CT aggregation plays in neuronal loss, its presence across a range of aging related dementia disorders indicates involvement in multi-proteinopathy driven neurodegeneration. To determine the TMEM CT aggregation propensity and neurodegenerative potential, we characterized a novel transgenic *C. elegans* model expressing the human TMEM CT fragment constituting the fibrillar core seen in FTLD cases. We found that pan-neuronal expression of human TMEM CT in *C. elegans* causes neuronal dysfunction as evidenced by behavioral analysis. Cytosolic aggregation of TMEM CT proteins accompanied the behavioral dysfunction driving neurodegeneration, as illustrated by loss of GABAergic neurons. To investigate the molecular mechanisms driving TMEM106B proteinopathy, we explored the impact of PGRN loss on the neurodegenerative effect of TMEM CT expression. To this end, we generated TMEM CT expressing *C. elegans* with loss of *pgrn-1*, the *C. elegans* ortholog of human PGRN. Neither full nor partial loss of *pgrn-1* altered the motor phenotype of our TMEM CT model suggesting TMEM CT aggregation occurs downstream of PGRN loss of function. We also tested the ability of genetic suppressors of tauopathy to rescue TMEM CT pathology. We found that genetic knockout of *spop-1, sut-2,* and *sut-6* resulted in weak to no rescue of proteinopathy phenotypes, indicating that the mechanistic drivers of TMEM106B proteinopathy may be distinct from tauopathy. Taken together, our data demonstrate that TMEM CT aggregation can kill neurons. Further, expression of TMEM CT in *C. elegans* neurons provides a useful model for the functional characterization of TMEM106B proteinopathy in neurodegenerative disease.

## Introduction: Impact of TMEM106B C-Terminal Fragment Aggregation on Neurodegenerative Disease

Diseases exhibiting accumulation of specific misfolded and aggregated proteins are generically referred to as proteinopathies and occur in association with neuronal degeneration and aging [1–4]. Protein aggregation of tau, amyloid beta peptide, α-synuclein, and TAR DNA binding protein 43 (TDP-43) represent well-characterized drivers of diverse proteinopathy disorders associated with dementia that include Alzheimer’s disease (AD), dementia with Lewy bodies (DLB), and frontotemporal lobar degeneration (FTLD) [5–7]. Accruing evidence suggest neurodegenerative events often involve co-occurring proteinopathies [8,9]. However, the molecular relationships between co-occurring pathological proteins remain incompletely understood. Improving interventions for proteinopathy disorders requires a better understanding of the interplay between distinct pathological protein species [10,11]. The transmembrane protein 106B (TMEM106B) has recently emerged as a new pathological protein implicated in multi-proteinopathies across a range of neurodegenerative conditions and in advanced aging [12–16].

TMEM106B is a single pass, lysosomal, type II transmembrane protein highly expressed in neurons and glial cells of the central nervous system (CNS) that plays an important role regulating lysosomal size, mobility, and acidity [17–20]. Decreased and increased expression of TMEM106B have been associated with severe lysosomal defects in neurons and oligodendrocytes [17,21–23]. TMEM106B was initially identified as a genetic risk factor for FTLD with TDP-43 (FTLD-TDP), with greater genetic impact on carriers of PGRN mutations [18,24–28]. Single nucleotide polymorphisms (SNPs) in TMEM106B have since been identified as risk modifiers for a variety of neurodegenerative diseases including AD, hippocampal sclerosis (HS), and Parkinson’s Disease (PD) [18,24,29–32].

With the understanding that lysosomal dysfunction occurs during neurodegeneration [33,34], determining the mechanistic role of lysosomal pathways in neuroprotection against proteinopathy remains unclear. Specifically, the molecular role TMEM106B plays in neurodegenerative disease requires further investigation. Several studies have explored the link between TMEM106B expression and neurodegenerative conditions, often with contradicting results. An analysis of TMEM106B SNPs in FTLD-TDP indicated that risk associated alleles increase protein levels through transcriptional activation [23]. Further, overexpression of TMEM106B in neurons was shown to increase lysosomal dysfunction caused by PGRN deficiency [35]. However, studies assessing the potential for TMEM106B reduction to ameliorate effects of PGRN loss in FTLD instead showed exacerbations in lysosomal dysfunction and FTLD related symptoms [21,36]. Different studies of TMEM106B risk alleles in AD cases reported both an increase and a decrease in TMEM106B expression in brain tissue [26,29].

Recently, due to advances in cryogenic electron microscopy (cryo-EM) previously unidentified, cytoplasmic protein aggregates in the brains of a diverse range of neurodegenerative diseases including FTLD, DLB, progressive supranuclear palsy (PSP), and PD were discovered to be homotypic aggregates of the C-terminal fragment of TMEM106B localized to the cytosol [12–16]. It is predicted that this C-terminal domain of TMEM106B is generally cleaved and degraded through normal lysosomal function [37]. Conditions of increased lysosomal burden may lead to accumulation and aggregation of this C-terminal protein in the cytoplasm, contributing to neurotoxicity in disease. In this study, we aim to model cytoplasmic TMEM106B C-terminal fragment (TMEM CT) aggregation *in vivo*, and assess the potential for this aggregation to impact neuronal degeneration in human disease.

Herein, we describe the first live organism transgenic model of TMEM106B C-terminal proteinopathy generated by expressing the TMEM106B C-terminal aggregation prone fragment constituting the fibrillar core in FTLD cases. To this end, we show that these TMEM106B fragments drive a suite of neurodegenerative phenotypes in *C. elegans* allowing genetic dissection of determinants of TMEM106B proteinopathy.

## Materials and Methods

### *C. elegans* strains and transgenics

**Table 1:**
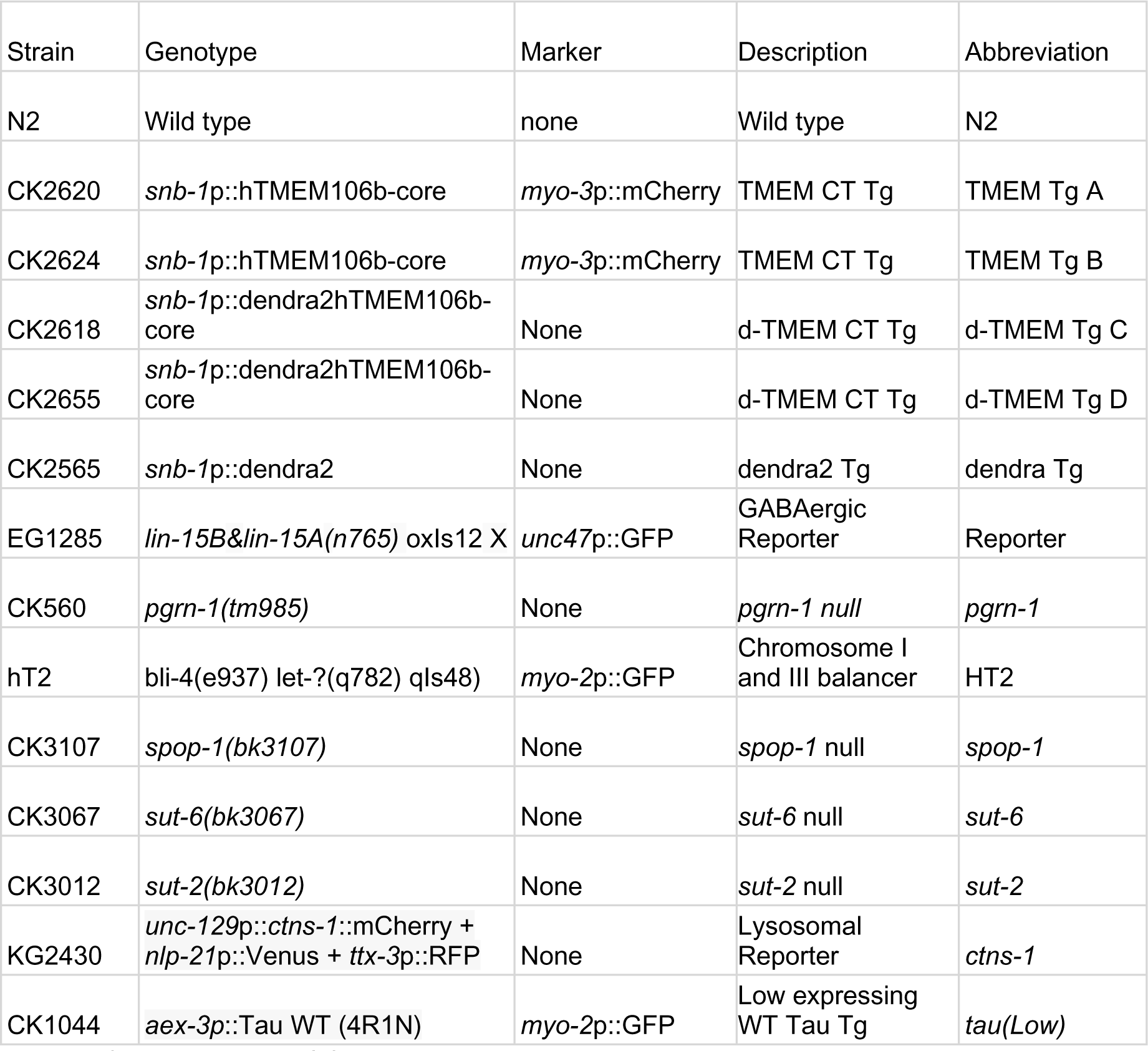
Strain List. List of *C. elegans* strains, genotypes, markers, descriptions, and abbreviations.

### Plasmids

To generate transgenes expressing the C-terminal fragment of human TMEM106B driven by a pan-neuronal snb-1 promoter, a *C. elegans* codon optimized DNA sequence encoding the C-terminal fibrillar core fragment of wild type TMEM106B (amino acids 120-254 of NP_001127704) was purchased Integrated DNA Technologies (IDT). The parental Psnb-1 vector was previously constructed [38] by inserting the snb-1 promoter sequences into the HindIII and BamHI sites of MCS I of plasmid pPD49.26 (a generous gift from Dr. A. Fire, Stanford University, Palo Alto, CA).

### Transgenics and strains

N2 (Bristol) was used as wild-type *C*. *elegans* and maintained as previously described [39]. The marker transgene *myo-3p*::mCherry was used at a concentration of 20 ng/μl as a co-injection marker as previously described [38]. The *snb-1p*::hTMEM106B-core and *snb-1p*::dendra2hTMEM106B-core transgenes described above were microinjected into N2 at a concentration of 150 ng/μl and 75 ng/μl respectively as previously described [40] to produce animals expressing TMEM106B and mCherry transgenes as extrachromosomal arrays. Extrachromosomal transgenes were integrated by exposing animals to UV radiation in a Stratalinker for 0.8 minutes. Progeny of irradiated animals were screened for 100% transmission of the transgene and integrated strains were backcrossed to N2 at least three times.

The *C. elegans* mutant *pgrn-1* (tm985) is a null mutation, featuring a 347 base pair deletion (13245/13246 – 13592/13593) in the pgrn-1 PGRN-like gene (obtained from the National BioResource Project, Japan) [41] was backcrossed to N2 five times as strain CK560.

Strain EG1285 has an integrated *unc-47p*::GFP transgene expressed in GABAergic neurons that clearly marks the cell bodies and axons of ventral cord VD and DD type inhibitory motor neurons [42]. *C*. *elegans* strains CK3107 (*spop-1* null), CK3067 (*sut-6* null), and CK3012 (*sut-2* null) were previously generated by CRISPR/cas9 mediated genome editing [43–45]. All TMEM106B transgenic strains described here contain integrated transgenes carrying human TMEM106B-core encoding cDNAs and a Pmyo-3::mCherry coinjection marker. CK2620 and CK2624 carry *snb-1p*::TMEM106B-core (TMEM Tg A and TMEM Tg B), while CK2618 and CK2655 carry *snb-1*p::dendra2hTMEM106b-core (d-TMEM Tg C and d-TMEM Tg D). Double mutants with TMEM106B expression were generated by crossing CK2620 or CK2624 with other strains carrying other transgenes or mutations of interest.

All strains were maintained at 20 °C on NGM plates seeded with OP50 *Escherichia coli* [39]. Genotypes were confirmed by PCR and sequencing for all strains with non-obvious phenotypes.

### Live *C*. *elegans* imaging

Live worms were mounted on a 4% agarose pad and immobilized with ∼50 mM sodium azide [Sigma] and covered with a glass slip. Representative images were acquired using a Nikon A1R confocal microscope with a 10x air objective or 40x or 100x oil immersion lens (Nikon USA, Melville, NY). Z-plane stacked images were flattened into a maximum intensity projection using ImageJ software.

### Immunohistochemistry

Day 1 adult worms were fixed in 1% formaldehyde solution and permeabilized by freeze cracking as described previously [46]. Fixed and permeabilized animals were counterstained with 300 nM 4,6-diamidino-2-phenylindole (DAPI) nuclear stain in PBS. Fixed whole animals were stained with anti-TMEM CTF monoclonal antibody (TMEM239) [47] at a dilution of 1:500. Alexa 488-conjugated anti-rabbit antibody (Invitrogen) and ALEXA 647-conjugated anti-mouse antibody was used as the secondary antibody at a dilution of 1:1000. 63 times magnification images were acquired by a Nikon A1R confocal microscope with a 100x oil immersion lens (Nikon USA, Melville, NY) or Andor Dragonfly 200 63x oil immersion lens (Andor UK, Bellfast) and processed in FIJI:imageJ.

### Behavioral analysis

NGM plates of day 1 adult *C*. *elegans* were flooded with 1 mL of M9 buffer (22 mM KH_2_PO_4_ monobasic, 42.3 mM Na_2_HPO_4_, 85.6 mM NaCl, 1 mM MgSO_4_), and swimming worms were pipetted onto a 35 mm unseeded NGM plate. Approximately 30 s following the addition of M9 buffer, worms were recorded swimming for 1 min at 14 frames per second. These videos were captured and analyzed with WormLab 2021 (MBF Bioscience). The frequency of body bends, or turns, as defined as a change in body angle of a least 20° from a straight line measured by the quarter points and midpoint of the worms, was quantified as a readout of locomotion. Worms tracked for less than 30 s were omitted from this analysis. At least three independent samples totaling at least 60 worms per strain were counted for every comparison [49].

### Neurodegeneration assays

Worms at larval stage 4 (L4) were selected and moved onto new NGM plates with OP-50. 24 hours later, live worms were placed on a 2% agarose pad containing 0.01% sodium azide to immobilize the worms. Worms were imaged under fluorescence microscopy and scored for number of GABAergic neurons. At least three independent samples totaling at least 60 worms per strain were counted for every comparison. Data were analyzed using GraphPad Prism software [50,51].

### Protein extraction

*C*. *elegans* was grown from hypochlorite-purified eggs at 20 °C for 3 days on 5XPEP plates until young adults. Worms were washed off plates with M9 buffer, collected by centrifugation, and grown further on egg plates as previously described [52]. Well-fed worms were collected from egg plates, harvested and isolated via sucrose floatation, washed into M9 buffer, and collected in 250 µL aliquots of packed worms prior to snap-freezing in liquid nitrogen to be stored at −70 °C. These worm samples were thawed, sonicated (11 x 15 s at 70% power and put on ice in between sonication steps) in TM-LS Buffer (10 mM Tris, 800 mM NaCl, 1 mM EDTA, 10% Sucrose, pH 7.5). Sarkosyl insoluble material was separated from soluble protein through ultra-centrifugation (2 spins at 140,000 x g for 30 mins each). The detergent insoluble material is not readily detectable without rigorous solubilization so a modified Aβ extraction protocol [53] was utilized to expose TMEM epitopes. Briefly, aggregated protein was resolubilized by resuspension and sonication in 90% formic acid (FA) to expose TMEM fibrillar epitopes. FA samples were dried in a speedvac to evaporate off formic acid. Dried disaggregated protein pellets were solubilized in 100 μL 5xSDS (46 mM Tris, 5 mM EDTA, 200 mM dithiothreitol, 50% sucrose, 5% sodium dodecyl sulfate, 0.05% bromophenol blue) protein sample buffer immediately after formic acid treatment. 200 μL aliquots were taken after initial solubilization in TM-LS buffer for soluble protein fractions and pelleted. Soluble protein was diluted with 60 μL 1× SDS protein sample buffer. Samples were sonicated at 70% for 15 s three times returning to ice between each replicate, boiled for 10 min at 95 °C, centrifuged at 13,000 × g for 2 min, and stored at −80 °C. Samples were loaded (6 μL) onto 4-15% precast criterion sodium dodecyl sulfate polyacrylamide gel electrophoresis gradient gels and transferred to PVDF per manufacturer’s protocol (Bio-Rad). The protein sizing ladder used was Precision Plus Protein Standards (Bio-Rad). The primary antibodies used were rabbit polyclonal anti-TMEM106B C-terminus antibody (TMEM 239) [15,47] at 1:800 and mouse anti-tubulin antibody E7 (Developmental Studies Hybridoma Bank) at 1:5,000. The secondary antibodies used were anti-mouse HRP or anti-rabbit HRP (Jackson Immuno Research) at 1:5,000. ECL substrate was used to visualize the membrane (Bio-Rad). Chemiluminescence signals were detected with the LiCor Imager and quantified with Fiji [54].

### Lifespan assay

L4 stage worms were picked onto NGM plates, and then transferred onto NGM plates with added 5-fluorodeoxyuridine (FUDR, 0.05mg/mL) to inhibit growth of progeny. Worms were housed at 25° C and scored every 1–2 days by gentle touching with a platinum wire. Failure to respond to touch was scored as dead. Worms that died as a result of FUDR or crawled off plates were censored [55]. Statistical analysis was performed using GraphPad Prism software.

## Results

### A transgenic *C. elegans* model of TMEM106B proteinopathy

The TMEM106B protein consists of an N-terminal cytosolic domain, a single transmembrane domain, and a C-terminal luminal domain. The C-terminal domain, from amino acids 120 – 255 consists of highly aggregate prone β-sheets, and makes up the fibril core recently discovered to aggregate in the brain across a number of neurodegenerative diseases (Figure 1A) [12–16]. *C. elegans* constructs expressing the human TMEM106B C-terminal fragment (TMEM CT) were designed to model the TMEM CT aggregation seen in a variety of neurodegenerative conditions. We generated transgenic *C*. *elegans* expressing human TMEM CT encoding cDNAs under control of a *C*. *elegans* pan-neuronal promoter) (Figure 1B) These transgenic strains are referred to as TMEM Tg A, and TMEM Tg B hereafter. We also generated transgenic *C. elegans* expressing dendra2 (a photo-convertible fluorescent protein) tagged human TMEM CT encoding cDNAs under control of the same pan-neuronal promoter (hereafter, d-TMEM CT--Figure 1C). These transgenic strains will be referred to as d-TMEM Tg C, and d-TMEM Tg D. To visualize TMEM CT expression, we conducted live imaging fluorescent microscopy experiments to observe the organismal location of TMEM CT accumulation. Representative images show expression of d-TMEM CT in d-TMEM Tg animals and dendra2 expression in dendra Tg animals, lacking the TMEM CT (Figure 2). Both the dendra Tg and the d-TMEM Tg strains contain cell body-filling puncta, primarily in the nerve ring and along the nerve cord. However, the d-TMEM Tg strain exhibits a noticeably lower degree of diffuse dendra2 signal, which suggests a high proportion of aggregated TMEM CT. Representative images of animals at larval stage 2 (L2) show a similar trend, indicating TMEM CT aggregation during worm development (Figure S1).

**Figure 1.**
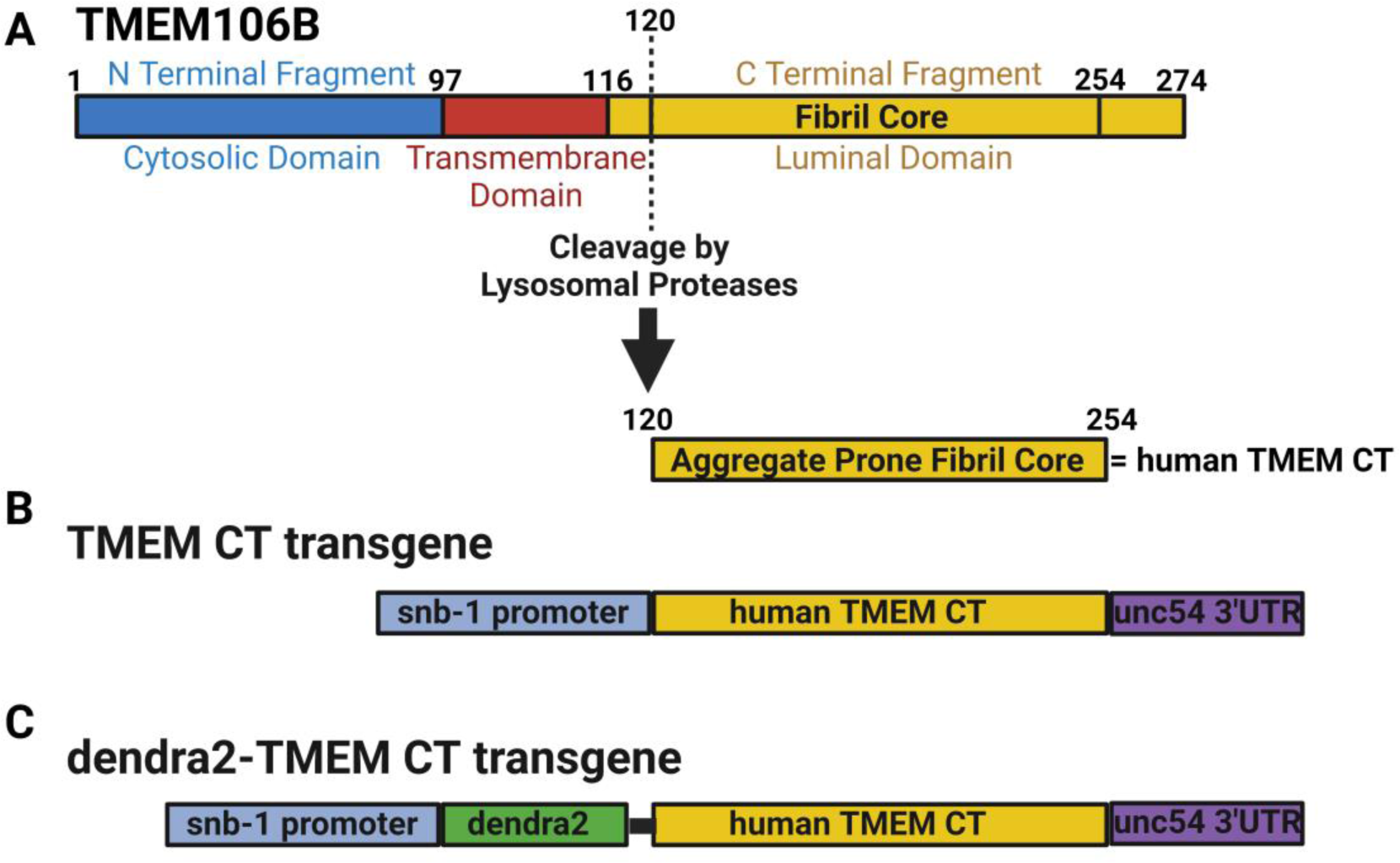
Transgenic insertion of TMEM CT in *C. elegans*. **A)** Transmembrane protein TMEM106B contains three domains: the N-terminal cytosolic domain, the transmembrane domain, and the C-terminal luminal domain. The luminal domain contains an aggregate prone fibril core (TMEM CT; AA 121-254). Diagram of transgenes containing **B)** TMEM CT or **C)** dendra2-tagged TMEM CT (d-TMEM CT) expressed under the pan-neuronal *snb-1* promoter. **D)** Experimental flow through for analysis of neuronal degeneration via thrashing assays and counts of GABAergic neurons by use of a fluorescent reporter strain.

**Figure 2.**
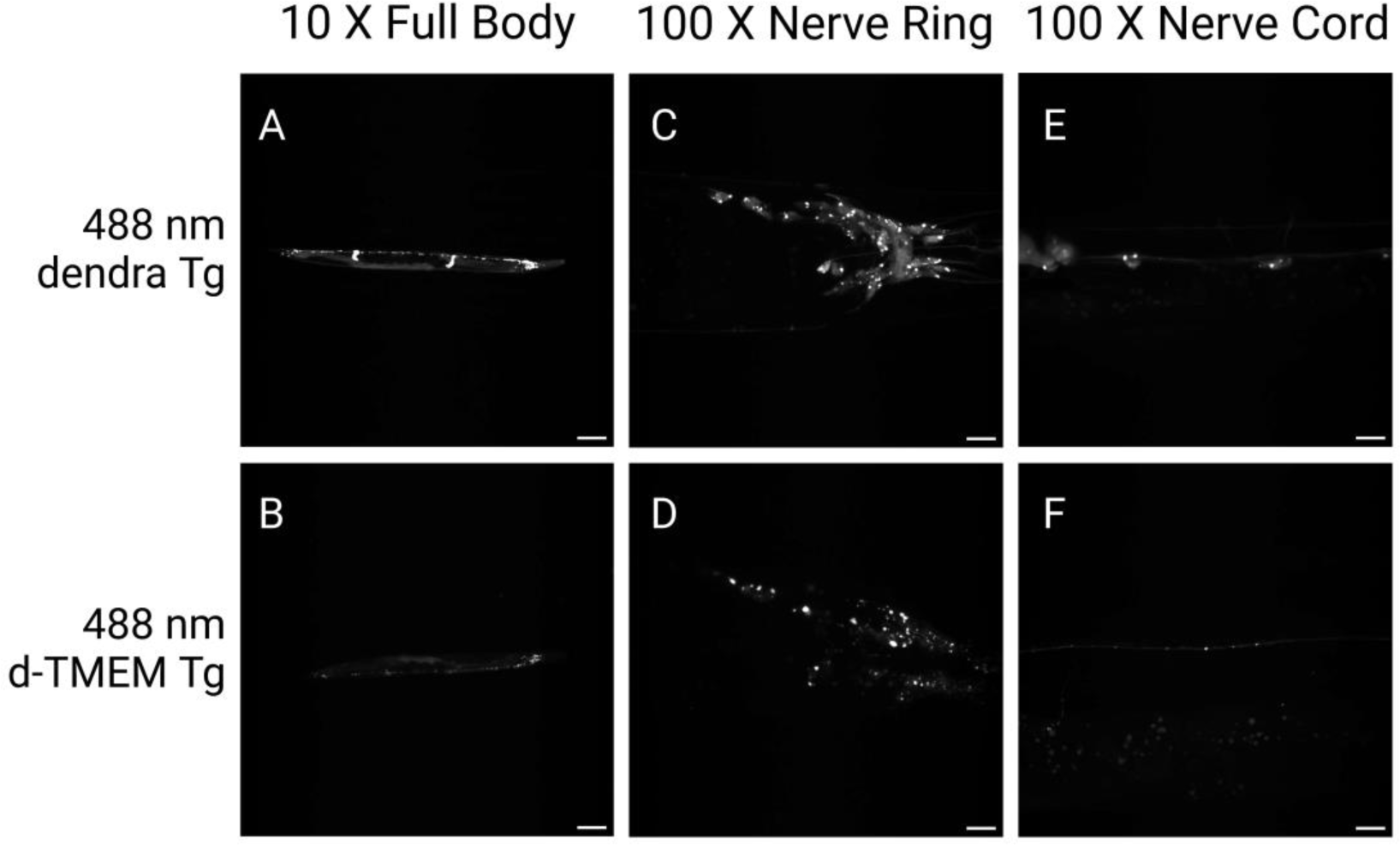
Transgenic *C*. *elegans* model expressing d-TMEM CT. 10X and 100X magnification images of day 1 adult dendra Tg and d-TMEM CT Tg *C*. *elegans*. **A-B)** 10X images of full worm body, signal indicating expression of the dendra2 fluorescent protein. Scale bar represents 100 µm. **C-D)** 100X images of nerve ring, signal indicating expression of the dendra2 fluorescent protein. Scale bar represents 10 µm. **E-F)** 100X images of nerve cord, signal indicating expression of the dendra2 fluorescent protein. Scale bar represents 10 µm.

In attempts to understand where in the cell TMEM CT aggregates are forming, we performed IHC of TMEM Tg strains, with DAPI staining for cell nuclei and an antibody against TMEM CT (TMEM 239) [47]. We observed TMEM CT protein accumulates in juxtanuclear aggregates (Figure 3 A-C). We also generated a cross between our d-TMEM Tg strains and a lysosomal reporter strain with the lysosomal protein, cystinosin homolog (CTNS-1), C-terminally fused to the red fluorescent protein, mCherry. Live imaging of these worms showed d-TMEM CT aggregates often formed adjacent to lysosomal signal, but did not appear to be engulfed by lysosomes (Figure 3 D-E).

**Figure 3.**
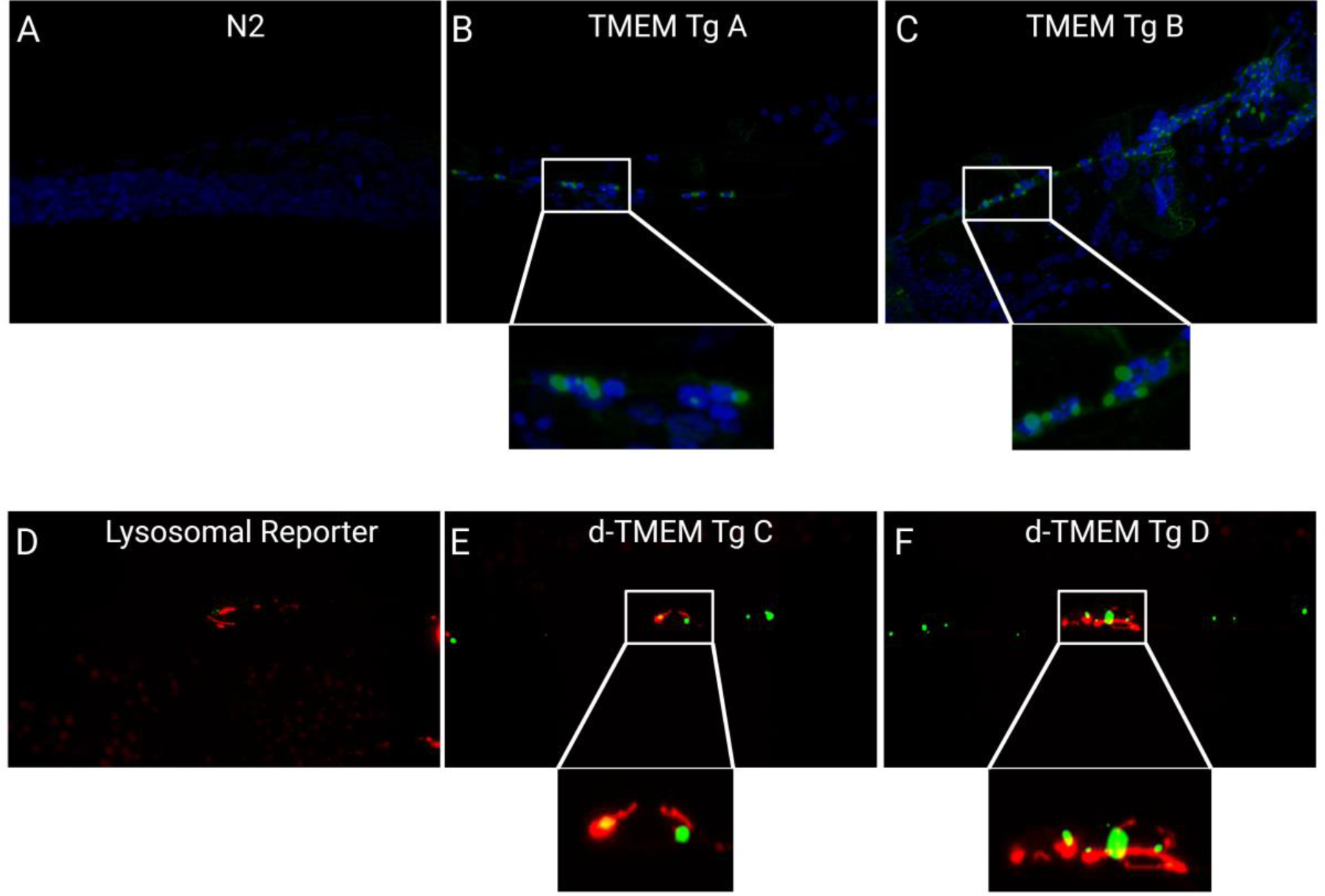
Localization of TMEM CT Aggregates. **(A-C)** 100X confocal images of TMEM CT immunofluorescence in day 1 adult N2 and TMEM CT Tg *C. elegans* indicate TMEM CT aggregates form juxta-nuclearly. Blue fluorescence indicates cell nuclei (DAPI), green fluorescence indicates TMEM CT (TMEM 239 rabbit antibody 1:500). **(D-E)** 100X confocal images of day 1 adult *ctns-1* and d-TMEM CT Tg *C. elegans* indicate TMEM CT aggregates form adjacent to, but not in lysosomes. Red fluorescence indicates CTNS-1::mCherry, and green fluorescence indicates d-TMEM CT. Images taken by Nikon Confocal Microscope and processed in FIJI:imageJ.

### TMEM CT aggregation drives neurodegeneration

Western blot analysis of transgenic *C*. *elegans* detected no TMEM CT in detergent soluble protein fractions (Figure 4A). However, based on extreme TMEM CT detergent insolubility in human disease, TMEM CT may accumulate in *C. elegans* as a highly detergent insoluble protein fragment. Studies with other highly aggregated amyloids have shown formic acid treatment to be an effective way of solubilizing detergent insoluble protein aggregates. Therefore we treated lysates from TMEM Tg animals with formic acid to expose aggregated TMEM106B protein epitopes [53]. Further western analysis of formic acid extracted protein aggregates confirmed TMEM CT expression (Figure 4A), and the evident aggregation propensity of this protein. We observed that d-TMEM Tg lines exhibited lower levels of TMEM CT expression and aggregation as compared to the TMEM Tg lines.

**Figure 4.**
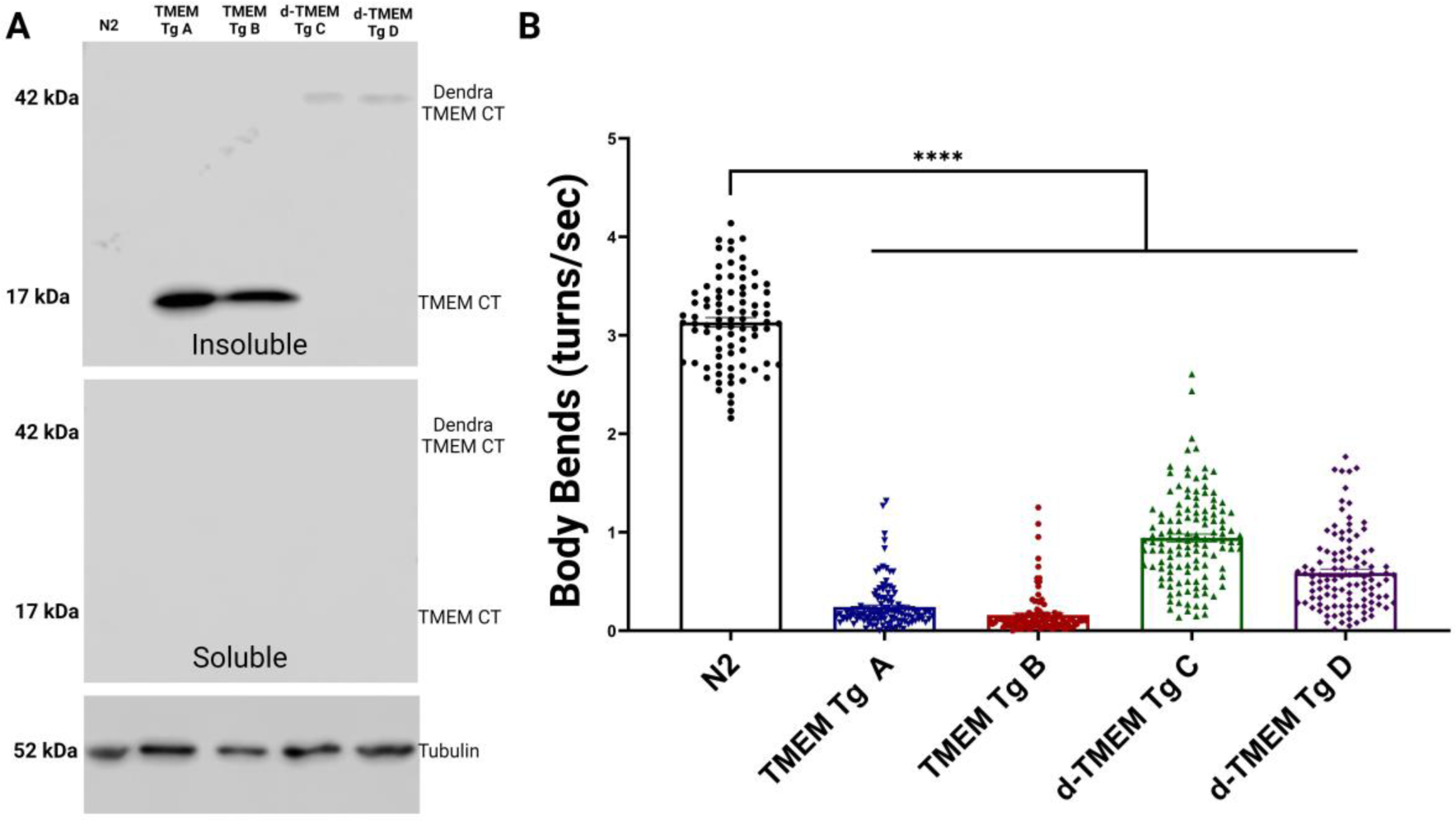
Aggregation of TMEM CT induces motor phenotype in Tg *C. elegans.* **A)** Western Blot analysis using TMEM CT specific antibody TMEM 239 depicts presence of TMEM CT (17kDa) and d-TMEM CT (42 kDa) in insoluble protein fractions of Tg *C. elegans* strains. No signal was observed in soluble protein fractions, confirming the highly aggregate prone nature of this protein fragment. **B)** Liquid thrashing assay of TMEM CT and d-TMEM CT strains assessed by computer analysis. n > 60, N = 3 for each strain. *C. elegans* expressing both TMEM CT and d-TMEM CT exhibit significantly impaired thrashing behavior as compared to N2 worms (p<0.0001), indicative of neuronal degeneration. P values denoted as **** for p<0.0001, error bars represent SEM.

Next, developmentally staged populations of TMEM CT transgenic *C*. *elegans* were assayed using motility (coordinated locomotion in liquid) as a sensitive readout of neuronal function. At day 1 of adulthood, TMEM CT and d-TMEM CT expressing lines performed significantly worse than the wild type N2 strain using the liquid thrashing assay as measured by observer independent digital video analysis (Figure 4B). This neuronal dysfunction was present at an early developmental timepoint, as TMEM and d-TMEM Tg strains were similarly impaired at the L2 stage of development (Figure S2). At both day one of adulthood, and stage L2 of development, the d-TMEM Tg strains exhibited a moderate degree of behavioral deficiency compared to the severe impairment in TMEM Tg animals. The abundance of TMEM CT appears related to drive the extent of behavioral deficits and suggests a tie between the abundance of TMEM CT aggregates and the severity of neuronal dysfunction.

To determine if behavioral deficits coincide with neuronal degeneration, we examined the integrity of GABAergic D-type motor neurons. We crossed TMEM CT lines with a reporter transgenic strain expressing GFP in GABAergic motor neurons under control of the *unc-47* promoter, allowing *in vivo* assessment of neurons in living TMEM Tg animals. At day 1 of adulthood, TMEM CT expressing strains showed significant loss of GABAergic motor neurons compared to the reporter strain, with TMEM Tg A losing around 1 of 19 neurons and TMEM Tg B losing close to 2 of 19 neurons on average (Figure 5). Given the lack of redundance within the *C. elegans* nervous system, we consider a ∼10% loss of GABAergic neurons to be a substantial. Similar analysis of TMEM Tg strains at the L2 stage of development indicated no detectable loss of GABAergic neurons during development (Figure S3). We take the observed neuronal dysfunction and lack of neuronal loss at the L2 stage as evidence that neuronal dysfunction precedes neuronal dysfunction.

**Figure 5.**
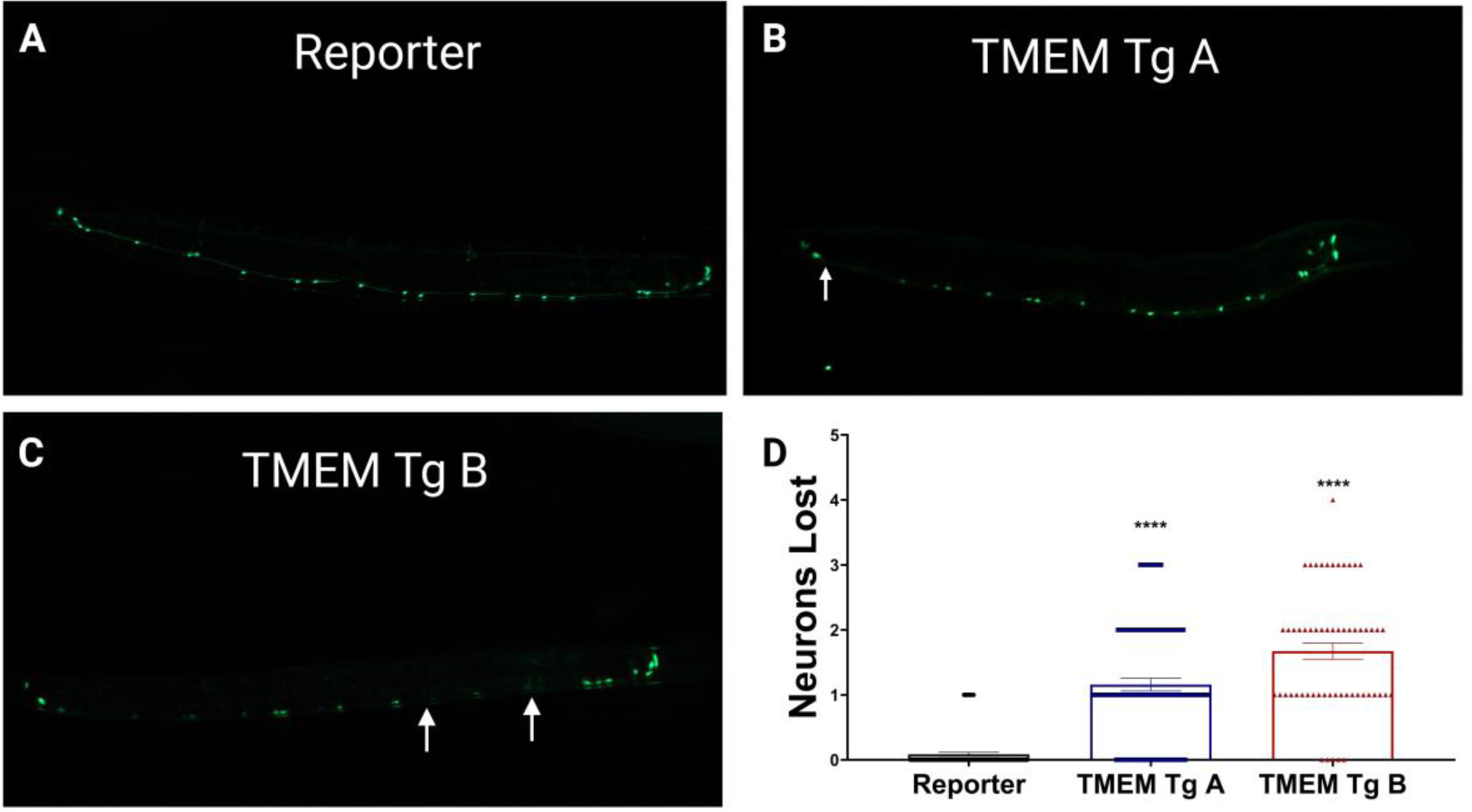
Neurotoxicity of TMEM CT in *C. elegans.* At day 1 of adulthood, TMEM Tg A loses on average 1 of 19 GABAergic neurons along the motor cord, while TMEM Tg B loses nearly 2 of 19 on average as visualized by the *unc-47*::GFP reporter (EG1285). Both Tg strains lose significantly more GABAergic neurons than the reporter strain (p<0.0001). **A-C)** Representative images for **A)** the reporter strain, **B)** TMEM Tg A, and **C)** TMEM Tg B. Arrows indicate lost neurons. **D)** Graphical representation of neuronal counts. n > 60, N = 3 for each strain. P values denoted as **** for p<0.0001, error bars represent SEM.

To assess the impact of TMEM CT aggregation on aging through adulthood, we conducted lifespan analysis on our transgenic strains. Under our conditions, the wild-type N2 strain, as well as *C. elegans* expressing dendra2 alone had a median survival of around 12 days of adulthood. Our TMEM CT Tg as well as d-TMEM CT Tg strains had median survivals ranging from 6 to 8 days of adulthood, representing a severe reduction in lifespan (Supplemental Figure S4 and Supplemental Table 1).

### Genetic interaction of TMEM Tg with Progranulin loss of function

SNPs in the human TMEM106B gene confer risk of development and severity for a number of neurodegenerative diseases, but most significantly in cases of FTLD-TDP caused by PGRN haplo-insufficiency [17,18]. Given that PGRN is a lysosomal protein governing protease activity, and partial loss of PGRN leads to impaired lysosomal function [56–58], we expect that loss of PGRN is involved in the mechanisms through which the TMEM CT ceases to be degraded and begins aggregation in disease.

To test this hypothesis, we crossed our TMEM Tg strains with *pgrn-1* null strains, and analyzed behavioral changes on TMEM Tg/*pgrn-1* (-/-) strains. We found that complete loss of *pgrn-1* had no significant impact on behavior of either TMEM Tg strain (Figure 6A). It is important to note that FTLD involves haplo-insufficiency of PGRN rather than a full knock out. A previous study showed that in *C. elegans*, complete loss of *pgrn-1* did not synergistically impact behavior of Tg TDP-43 mutant strains, while heterozygous loss of *pgrn-1* significantly impaired behavior in the same Tg strain [56].

**Figure 6.**
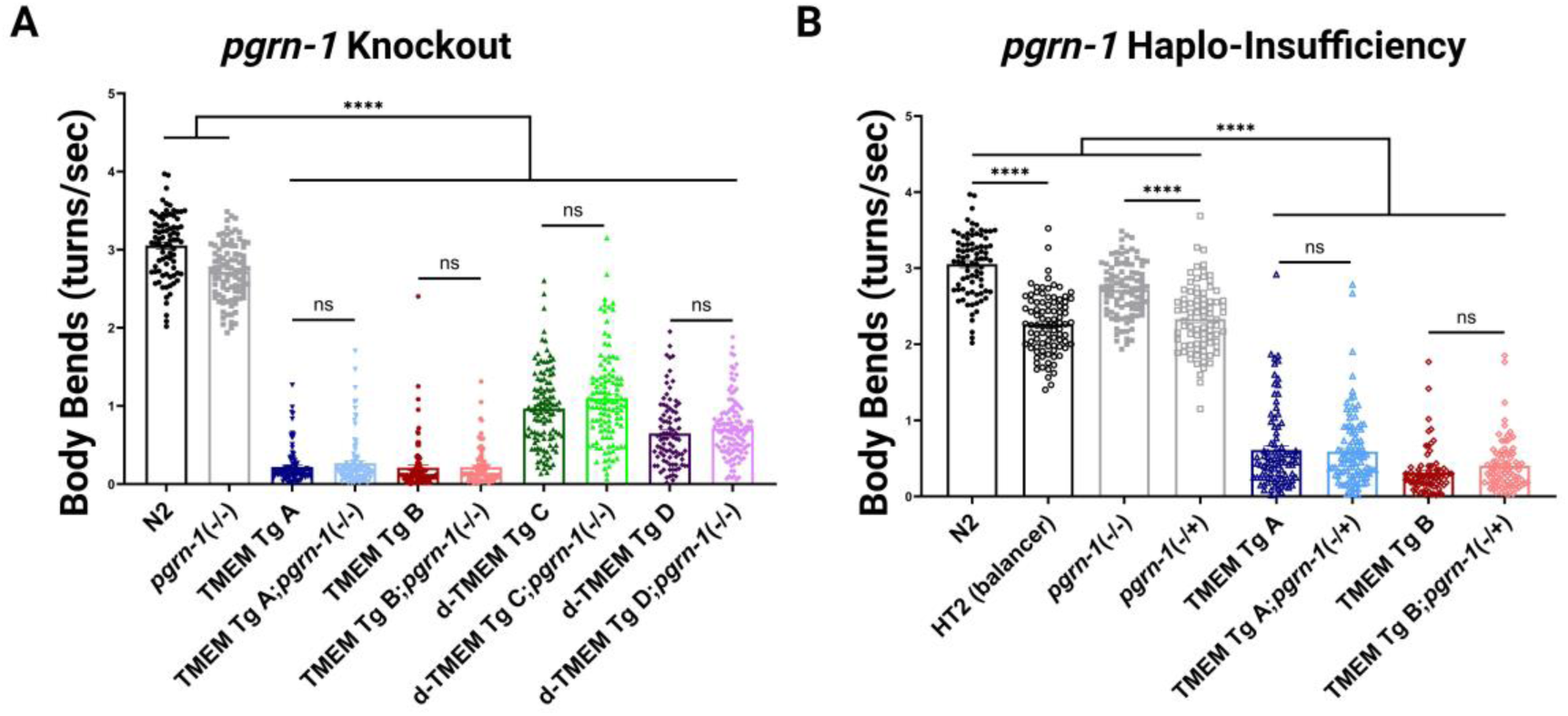
Loss of *pgrn-1* Does Not Impact Behavior of TMEM CT *C. elegans*. **A)** Liquid thrashing assay of TMEM CT expressing strains with complete loss of *pgrn-1* assessed by computer analysis. n > 60, N = 3 for each strain. Complete loss of *pgrn-1* had no significant impact on worm behavior. **B)** Liquid thrashing assay of TMEM CT strains with partial loss of *pgrn-1* assessed by computer analysis. n > 60, N = 3 for each strain. Similarly, partial loss of *pgrn-1* had no significant impact on worm behavior. P-values noted as ns for no significance or **** for p<0.0001, error bars represent SEM.

We generated *pgrn-1* heterozygotes by crossing *pgrn-1* null and TMEM Tg animals with the balancer hT2, a homozygous inviable balancer for *C. elegans* chromosomes I and III. Unlike the impact of heterozygous *pgrn-1* on behavior in a Tg TDP-43 model, we observed no significant effect of *pgrn-1* haplo-insufficiency on TMEM Tg behavior with TMEM Tg;*pgrn-1*(-/+) worms performing similarly to TMEM Tg strains (Figure 6B).

### Interactions of TMEM with common suppressors of tau pathology

TMEM106B pathology co-occurs with tau pathology in PSP, PD, FTLD-tau, and AD [12–16]. In addition, TMEM106B mutation or loss of function can influence tau pathology in transgenic mice [59,60]. Our previous studies using forward genetic screens to isolate suppressors of tau pathology in tau Tg *C*. *elegans* have revealed a number of modifier genes, including *spop-1*, *sut-2*, and *sut-6* [43,44,49,61–64]. To determine whether there are shared mechanisms of neurotoxicity between tau and TMEM CT, we tested whether loss of *spop-1, sut-2,* or *sut-6* might have an effect on the phenotype of our TMEM Tg model.

TMEM Tg A; *spop-1* and TMEM Tg B; *spop-1* performed significantly better on the thrashing assay than TMEM Tg A and TMEM Tg B alone (Figure 7A). However, with only a 23.6 and 9.8% rescue of phenotype respectively, we do not consider *spop-1* a strong suppressor of TMEM CT. Similarly, TMEM Tg A;*sut-6* and TMEM Tg B; *sut-6* performed significantly better than TMEM Tg A and TMEM Tg B (Figure 7B), but only showed a 26.8 and 8.4% return of function respectively. Lastly, loss of *sut-2* did not result in any significant modification of phenotype for either TMEM Tg strain (Figure 7C). For comparison, loss of function mutations in *spop-1, sut-6, and sut-2* rescue tau Tg *C. elegans* models of tauopathy from 50 – 100% of non-Tg motility levels [43,44,61].

**Figure 7.**
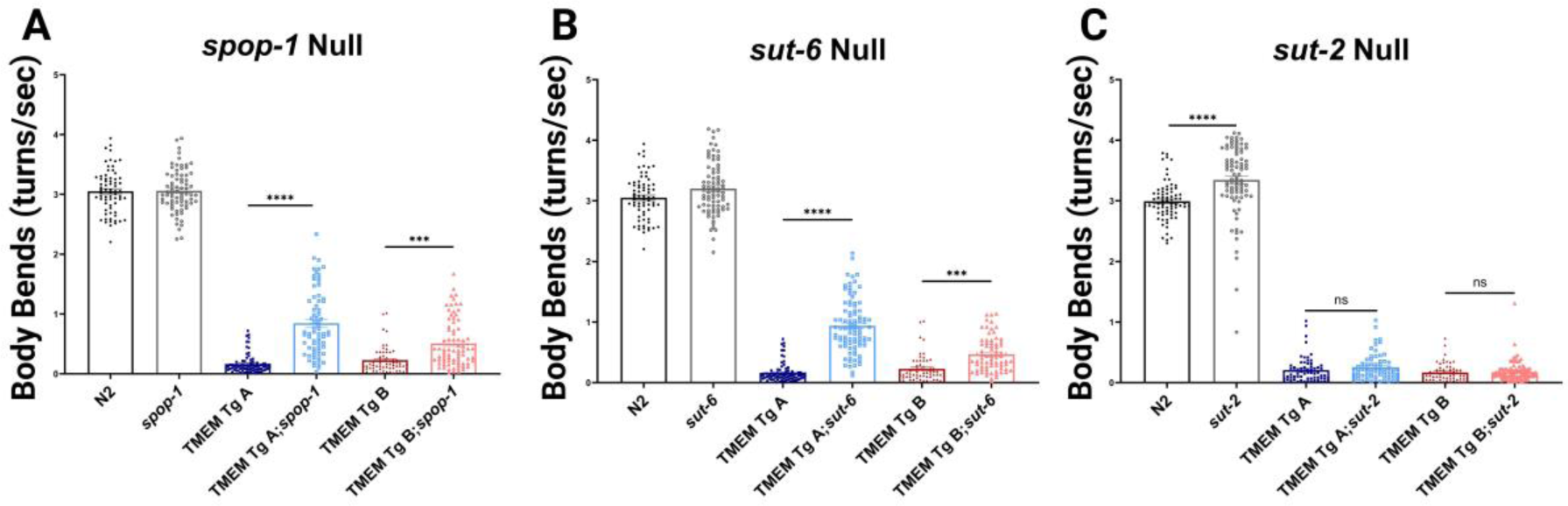
Suppressors of Tau and TDP-43 Pathology Only Weakly Modify TMEM CT Behavioral Deficit. **A)** Liquid thrashing assay of TMEM CT expressing strains with complete loss of SPOP-1 assessed by computer analysis. n > 60, N = 3 for each strain. Loss of *spop-1* confers a weak, but statistically significant return of function for TMEM CT expressing strains. **B)** Liquid thrashing assay of TMEM Tg strains with complete loss of *sut-6* assessed by computer analysis. n > 60, N = 3 for each strain. Loss of *sut-6* confers a weak, but statistically significant return of function for TMEM CT expressing strains. **C)** Liquid thrashing assay of TMEM Tg strains with complete loss of *sut-2* assessed by computer analysis. n > 60, N = 3 for each strain. Loss of *sut-2* had not significant effect on mobility of TMEM CT expressing strains. P values noted as *** for p<0.0001, error bars represent SEM.

## Discussion

Over the past decade SNPs in TMEM106B have emerged as a reproducible genetic risk factor for development and severity of neurodegenerative diseases, especially FTLD-TDP [17–19,21–23,25–30]. Previous studies exploring the role of TMEM106B in neurodegeneration have focused primarily on overall levels of protein expression or its role in lysosomal function. TMEM106B expression can be reduced in the brains of AD patients [26], and some studies have shown that loss of TMEM106B can induce severe lysosomal abnormalities in neurodegenerative models, especially in the context of PGRN loss [21,36,59]. Conversely, TMEM106B reduction has rescues lysosomal function caused by deletion of PGRN [28,65] and risk associated TMEM106B SNPs may alter chromatin architecture, increasing TMEM106B expression, and induce toxic lysosomal defects [23,32,35]. Due to the conflicting nature of these findings, the role of TMEM106B in neurodegeneration remains unclear. Recently, the C-terminal fragment of TMEM106B was demonstrated to form amyloid fibrils in the brains of patients across a diverse range of neurodegenerative diseases including AD and many ADRDs [12–14,16,66]. Furthermore, aggregation of this protein fragment has been observed in the brain naturally with age [15]. While the relative toxicity of the accumulating TMEM106B CT aggregates remains uncertain, the presence of TMEM106B amyloid across a variety of neurodegenerative diseases indicates involvement in multi-proteinopathy driven neurodegeneration. Understanding the aggregation propensity of TMEM106B as well as the neurodegenerative potential of TMEM106B fibrillization requires new animal and cellular models of the TMEM106B proteinopathy occurring in aging related neurodegenerative disease.

In this study, we describe a series of new transgenic *C. elegans* lines used to model TMEM106B proteinopathy by expressing TMEM CTs in all neurons. We highlight this work represents the first live organism model of TMEM CT aggregation in neurons, and recapitulates key aspects of TMEM CT associated neurodegenerative change. The TMEM106B transgene expresses a 135 amino acid C-terminal fragment present in the fibrillar core of TMEM106B amyloid that is predicted to have a high β-sheet strand content with high aggregation propensity [12]. Western blot analysis of TMEM106B transgenic lines confirmed that when the TMEM CT is expressed in *C elegans,* the protein fragment can only be detected in the detergent insoluble fraction. This, combined with the requirement for rigorous aggregate solubilization using formic acid to permit TMEM CT detection speaks to the highly aggregated state of TMEM106B expressed in *C. elegans*. The *C. elegans* genome lacksa homolog of TMEM106B, and as such we cannot predict whether full length TMEM106B would be processed into TMEM CT fragments in the authentic mammalian manner when expressed in *C. elegans* neurons. One common post translational modification of TMEM106B, including on the C-terminal domain, is glycosylation [16]. The expected sizes of our expressed protein fragments are 17kDa for the C-terminal fragment alone, and 42 kDa for dendra2 fused to the TMEM CT. Based on the 17 kDa size of the TMEM CT and 42 kDa size of d-TMEM CT on Western blot, these protein fractions did not appear to be glycosylated in our model, although the necessity of formic acid extraction to solubilize TMEM106B CTF aggregates may mask the endogenous glycosylation state of the proteins. Behavioral analysis of day 1 TMEM Tg adult animals revealed profound behavioral abnormalities indicative of neuronal dysfunction and possible neurodegeneration. This was corroborated by counts of GABAergic motor neurons, which revealed TMEM CT Tg strains exhibited consistent loss of GABAergic neurons. While loss of an average 2 neurons per animal at day 1 of adulthood may seem minor, it represents a loss of more than 10% of this non-redundant neuronal population consistent with the manifestation of behavioral phenotypes. Lifespan analysis of TMEM CT Tg strains indicated a severe decline in lifespan as a result of TMEM CT aggregation. Taken together these data clearly demonstrate the high aggregation propensity of the TMEM106B CTF and support a strongly neurotoxic character of these aggregates. Furthermore, analysis of transgenic *C. elegans* at the L2 stage of development show TMEM CT aggregation along with evidence of neuronal dysfunction that precedes loss of neurons, as no GABAergic neurons were lost at the L2 stage. Thus, the C-terminal domain of TMEM106B should be considered a pathogenic protein, and the concept of TMEM106B proteinopathy warrants further analysis both in the context of age-related changes to TMEM106B and multi-proteinopathy disorders.

To explore potential genetic interactions between TMEM106B and PGRN in *C. elegans*, we explored the consequences of *pgrn-1* loss of function on TMEM CT proteinopathy related phenotypes. TMEM106B SNPs have been seen to have the most significant impacts on risk and disease severity in cases involving PGRN haplo-insufficiency [17–19], and alterations in TMEM106B expression can exacerbate lysosomal abnormalities caused by loss of PGRN [17–19,21,25,28,35,36,65]. We expect lysosomal dysfunction induced by loss of PGRN impairs TMEM106B processing, resulting in aggregation and TMEM106B proteinopathy. In order to test this hypothesis, we generated TMEM Tg strains with either full or partial loss of *pgrn-1* and assessed behavioral readouts. We saw no evidence of *pgrn-1* modification of behavioral abnormalities induced by TMEM106B proteinopathy caused by either full of partial loss of *pgrn-1*, indicating no interaction between *pgrn-1* and TMEM106B in this model system. We attribute this negative finding to the fact that all detectable TMEM CT expressed appears to be aggregated. If loss of PGRN instigates TMEM106B proteinopathy by impairing the lysosome’s ability to process this protein fragment, our model expressing a highly aggregation prone fragment bypasses the need for altered lysosomal processing by PGRN loss of function. This would result in no observed alteration in TMEM proteinopathy related phenotypes. Mammalian animal model systems may be helpful in fully recapitulating the influence of PGRN loss of function on TMEM106B processing.

In our previous work on proteinopathy disorders using *C. elegans* we have uncovered potent genetic modifiers of tau proteinopathy including SPOP-1, SUT-2/MSUT-2, and SUT-6/NIPP1. Genetic loss of function in these genes strongly suppresses the tauopathy phenotypes evident in transgenic *C. elegans* [43,44,49,61–64,67]. Consequently, we investigated whether loss of these strong regulators of tau proteinopathy might impact pathological TMEM106B. Our analysis revealed only modest rescue of TMEM106B proteinopathy phenotypes by *spop-1* or *sut-6* loss of function. Furthermore, *sut-2* loss of function exhibited no rescue of TMEM106B proteinopathy. With little to no rescue, we do not consider any of these genes as effective suppressors of TMEM106B pathology. This may suggest that TMEM106B proteinopathy involves unique mechanisms of neurotoxicity as compared to tau.

Emerging evidence indicates multi-proteinopathy appears to be a much more common driver of neurodegeneration than previously thought, and understanding how each proteinopathy interacts with the others will prove crucial for developing targeted therapeutic strategies [8–11,68]. Having shown the neuron killing propensity of TMEM106B CT aggregates, exploring the interplay between TMEM106B and other common proteinopathies appears the logical next step in investigating the molecular basis of multiple etiology dementia.

## List of Abbreviations

Transmembrane Protein 106B (TMEM106B)

Frontotemporal Lobar Degeneration (FTLD)

Progranulin (PGRN)

Cryo-electron microscopy (Cryo-EM)

TMEM106B C-terminal fragment (TMEM CT)

Progressive Supranuclear Palsy (PSP)

Dementia with Lewy Bodies (DLB)

TAR DNA binding protein 43 (TDP-43)

Alzheimer’s Disease (AD)

Central nervous system (CNS)

FTLD with TDP-43 pathology (FTLD-TDP)

Single nucleotide polymorphism (SNP)

Hippocampal Sclerosis (HS)

Parkinson’s Disease (PD)

Formic Acid (FA)

Phosphorylated Tau (p-Tau)

Larval Stage 2 (L2)

Larval Stage 4 (L4)

## Declarations

### Ethics approval and consent to participate

This study has no human or mammalian data.

### Consent for publication

All authors approve of this manuscript and consent to publication.

### Competing interests

The authors have no competing interests to declare.

## Availability of Data and Materials

All data supporting the conclusions of this article are included within the article and in additional files provided.

## Funding

This work was supported by grants from the Department of Veterans Affairs [I01BX004044 to N.L & IK6BX006467 to B.K.] and National Institutes of Health [R01AG066729 to N.L. & R01NS064131 to B.K.].

## Author’s Contributions

R.R. performed experiments, analyzed data, edited the manuscript and wrote the first draft manuscript. A.S performed experiments, provided methodology, generated strains, and editing the manuscript. P.J.M. performed experiments and edited the manuscript. R.L.K. provided methodology, analyzed data and edited the manuscript. N.FL. provided methodology, analyzed data, and edited the manuscript. B.C.K. conceived and oversaw the study, analyzed data, and wrote the manuscript.

## Acknowledgements

We thank the reviewers for helpful comments and suggestions. We thank Heather Currey, Elaine Loomis, Isaac Miller, Jade Stair, Ashley Sciocchetti, Brandon Henderson, Asia Beale, Lisa Chang, and Kailee Parrott for outstanding technical assistance. We thank Sjors Scheres for generously sharing the highly specific TMEM239 antibody and the Developmental Studies Hybridoma Bank (NICHD) for the β-tubulin antibody E7. Some strains were provided by the CGC, which is funded by NIH Office of Research Infrastructure Programs (P40 OD010440), and the National Bioresource Project (Japan). Illustrations Created with Biorender.

## Supplementary Figures

**Supplemental Figure S1.**
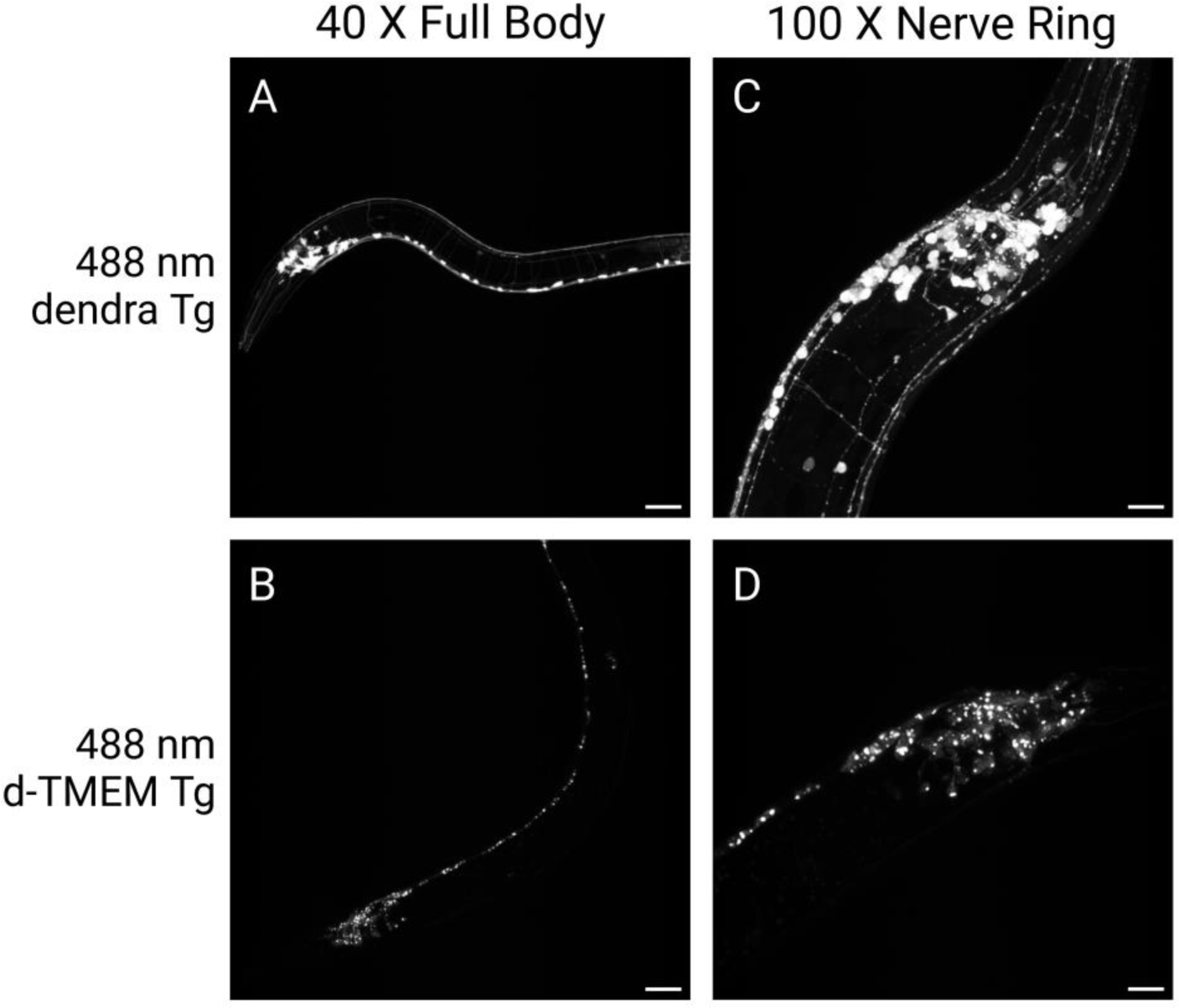
Transgenic *C*. *elegans* model expressing d-TMEM CT at L2 stage of development. 40X and 100X magnification images of L2 staged dendra Tg and d-TMEM CT Tg *C*. *elegans*. **A-B)** 40X images of full worm body, signal indicating expression of the dendra2 fluorescent protein. Scale bar represents 25 µm. **C-D)** 100X images of nerve ring, signal indicating expression of the dendra2 fluorescent protein. Scale bar represents 10 µm.

**Supplemental Figure S2.**
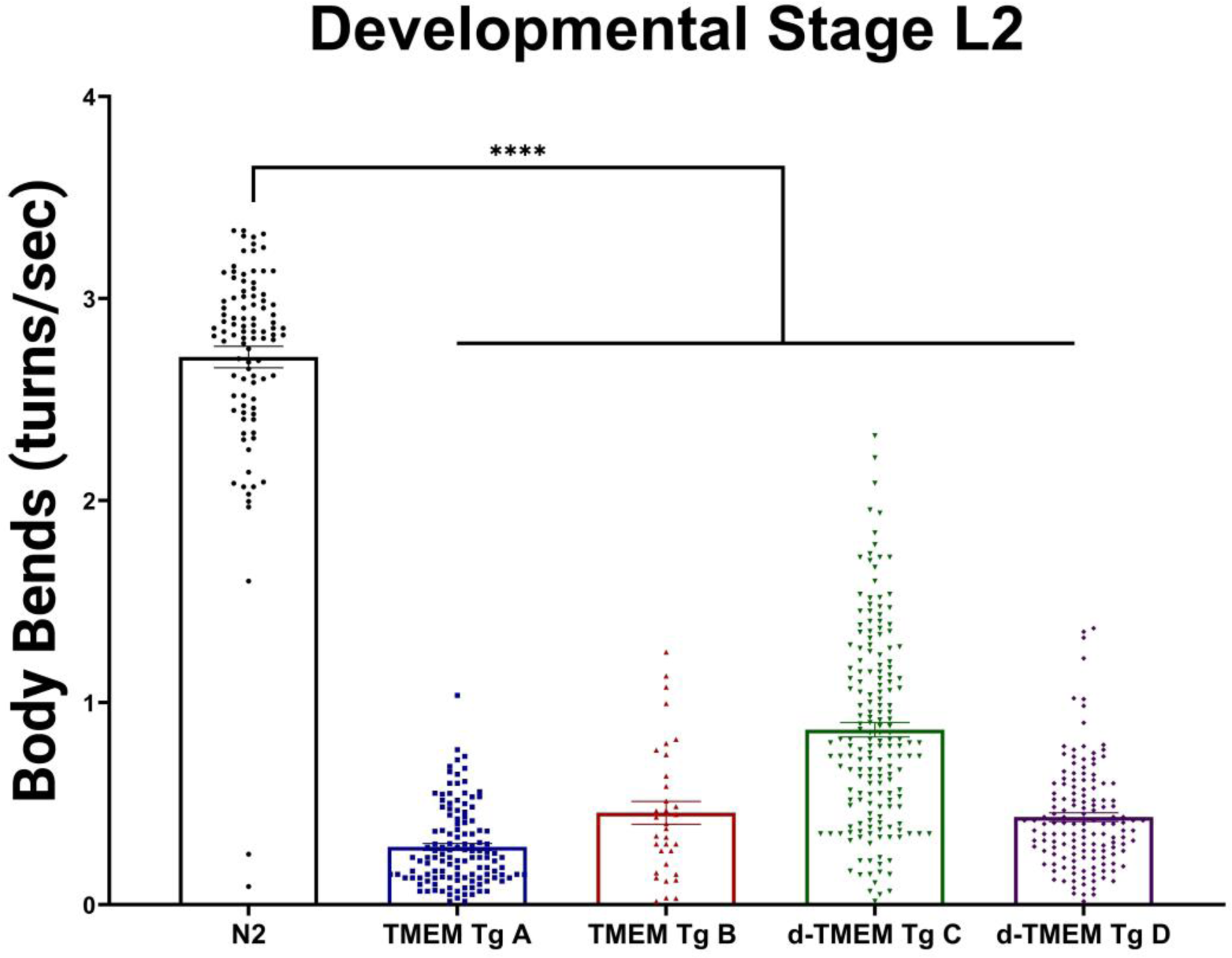
TMEM CT expression impairs locomotion at the L2 stage of development. Liquid thrashing assay of TMEM CT and d-TMEM CT strains assessed by computer analysis, mean of replicate averages. n > 34, N = 3 for each strain. *C. elegans* expressing both TMEM CT and d-TMEM CT exhibit significantly impaired thrashing behavior as compared to N2 worms (p<0.0001), indicative of neuronal degeneration. P values denoted as **** for p<0.0001, error bars represent SEM.

**Supplemental Figure S3.**
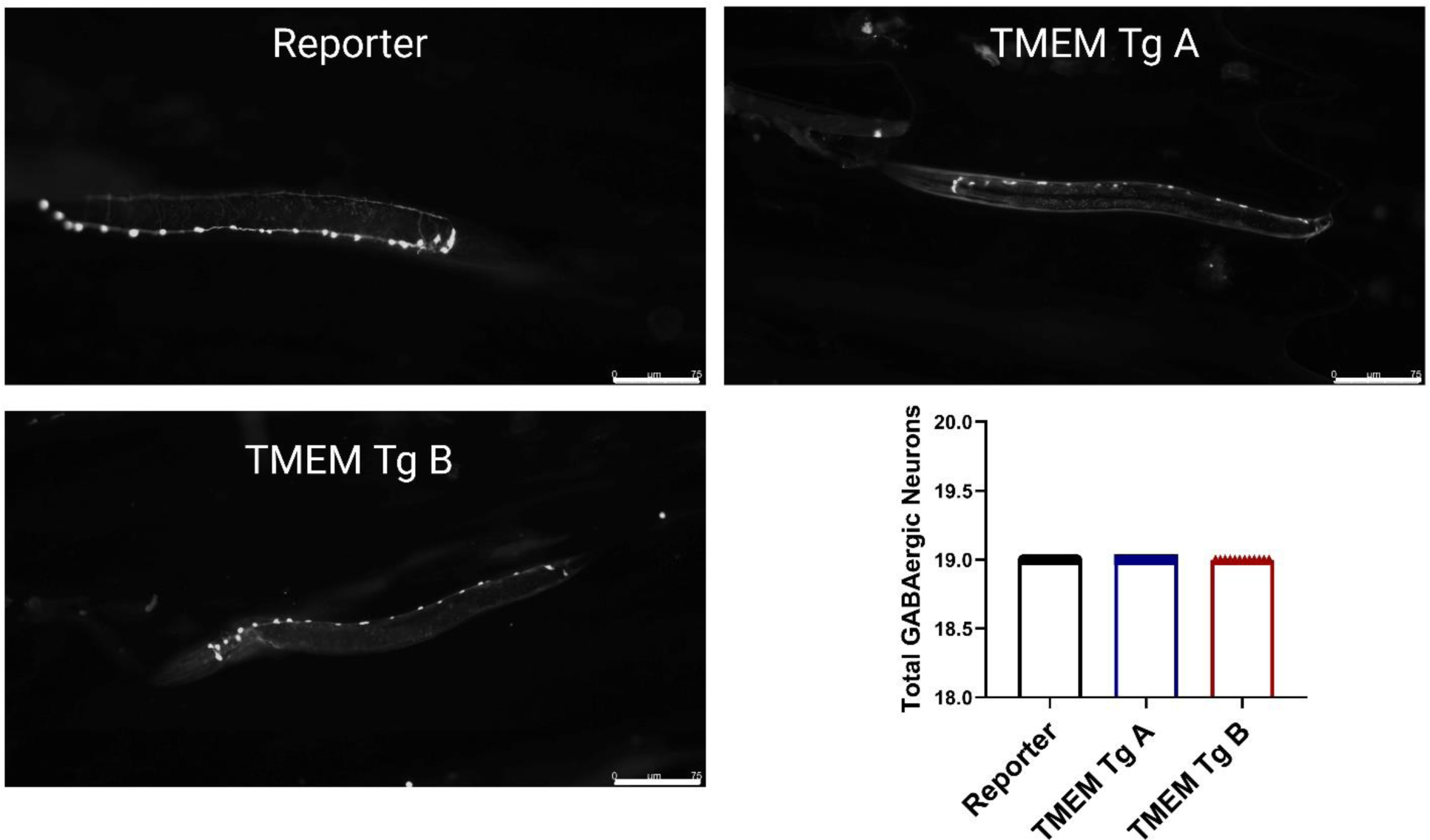
TMEM CT Aggregation Does Not Induce Neuronal Loss in L2 Stage *C. elegans*. At the L2 stage of development, neither TMEM Tg A nor TMEM Tg B strain lose any of their 19 GABAergic neurons as visualized by the *unc-47*::GFP reporter (EG1285). Representative images for the reporter strain, TMEM Tg A, and TMEM Tg B and graphical representation of neuronal counts. n > 12 for each strain. Scale bar indicates 75 µM, and error bars represent SEM.

**Supplemental Figure S4.**
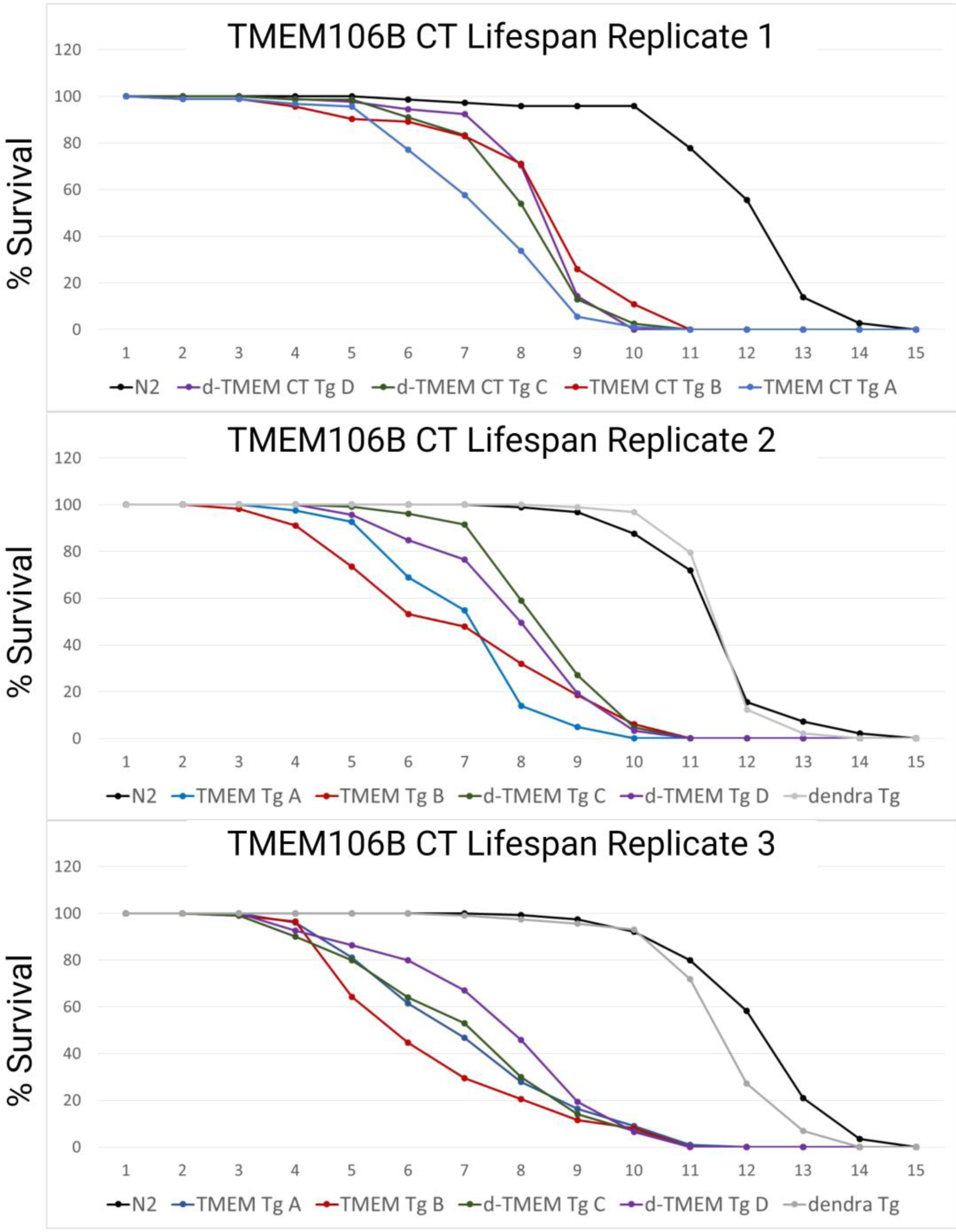
TMEM CT Aggregation Severely Decreases Lifespan of *C. elegans.* Lifespan assays of transgenic and wildtype *C. elegans* under FUDR treatment at 25° C. n > 72 worms per strain, N=3. Wild type (N2) worms and transgenic worms expressing only dendra2 had a median survival of around 12 days of adulthood. TMEM CT Tg and d-TMEM CT Tg strains had a significantly reduced median survival ranging from 6 to 8 days of adulthood.

**Supplemental Table S1:**
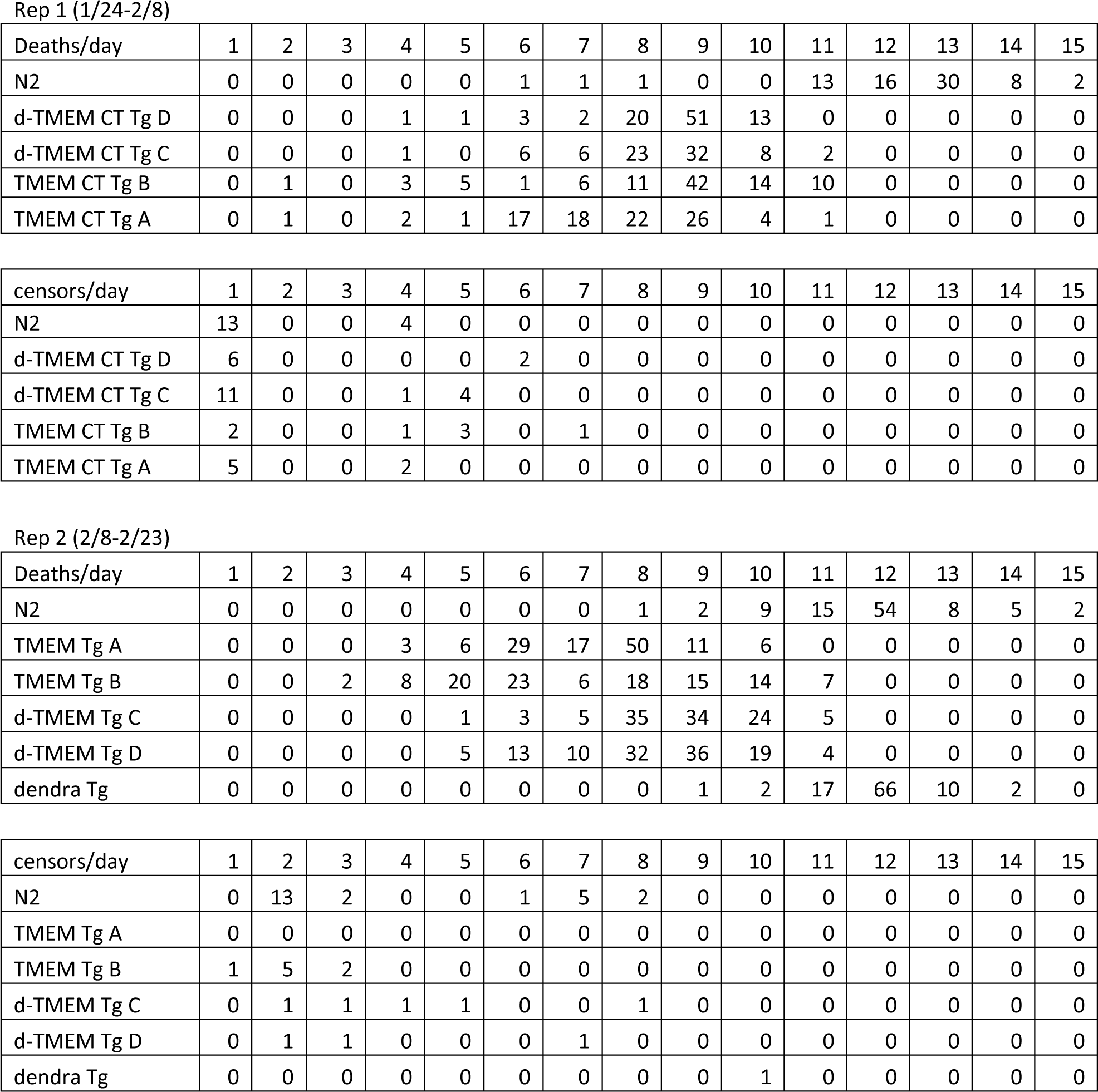
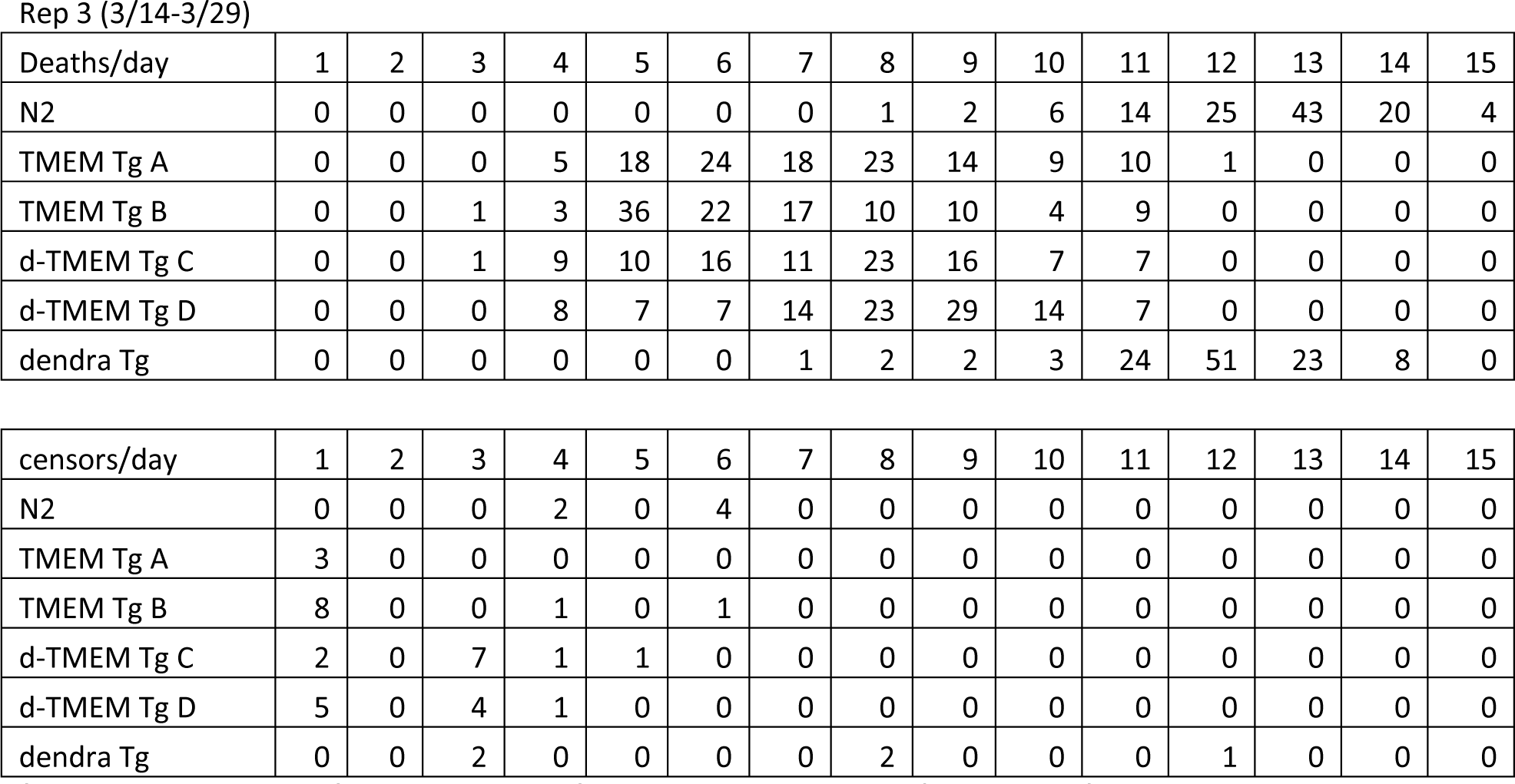
Deaths and Censures per Day For *C. elegans* Strains During Lifespan Assay. Worms counted as dead when one did not respond to gentle touch from platinum wire. Worms that burst from FUDR treatment or worms that crawled off of the plate during assay were censored.

| Deaths/day | 1 | 2 | 3 | 4 | 5 | 6 | 7 | 8 | 9 | 10 | 11 | 12 | 13 | 14 | 15 |
| --- | --- | --- | --- | --- | --- | --- | --- | --- | --- | --- | --- | --- | --- | --- | --- |
| N2 | 0 | 0 | 0 | 0 | 0 | 1 | 1 | 1 | 0 | 0 | 13 | 16 | 30 | 8 | 2 |
| d-TMEM CT Tg D | 0 | 0 | 0 | 1 | 1 | 3 | 2 | 20 | 51 | 13 | 0 | 0 | 0 | 0 | 0 |
| d-TMEM CT Tg C | 0 | 0 | 0 | 1 | 0 | 6 | 6 | 23 | 32 | 8 | 2 | 0 | 0 | 0 | 0 |
| TMEM CT Tg B | 0 | 1 | 0 | 3 | 5 | 1 | 6 | 11 | 42 | 14 | 10 | 0 | 0 | 0 | 0 |
| TMEM CT Tg A | 0 | 1 | 0 | 2 | 1 | 17 | 18 | 22 | 26 | 4 | 1 | 0 | 0 | 0 | 0 |

| censors/day | 1 | 2 | 3 | 4 | 5 | 6 | 7 | 8 | 9 | 10 | 11 | 12 | 13 | 14 | 15 |
| --- | --- | --- | --- | --- | --- | --- | --- | --- | --- | --- | --- | --- | --- | --- | --- |
| N2 | 13 | 0 | 0 | 4 | 0 | 0 | 0 | 0 | 0 | 0 | 0 | 0 | 0 | 0 | 0 |
| d-TMEM CT Tg D | 6 | 0 | 0 | 0 | 0 | 2 | 0 | 0 | 0 | 0 | 0 | 0 | 0 | 0 | 0 |
| d-TMEM CT Tg C | 11 | 0 | 0 | 1 | 4 | 0 | 0 | 0 | 0 | 0 | 0 | 0 | 0 | 0 | 0 |
| TMEM CT Tg B | 2 | 0 | 0 | 1 | 3 | 0 | 1 | 0 | 0 | 0 | 0 | 0 | 0 | 0 | 0 |
| TMEM CT Tg A | 5 | 0 | 0 | 2 | 0 | 0 | 0 | 0 | 0 | 0 | 0 | 0 | 0 | 0 | 0 |

Rep 2 (2/8-2/23)
| Deaths/day | 1 | 2 | 3 | 4 | 5 | 6 | 7 | 8 | 9 | 10 | 11 | 12 | 13 | 14 | 15 |
| --- | --- | --- | --- | --- | --- | --- | --- | --- | --- | --- | --- | --- | --- | --- | --- |
| N2 | 0 | 0 | 0 | 0 | 0 | 0 | 0 | 1 | 2 | 9 | 15 | 54 | 8 | 5 | 2 |
| TMEM Tg A | 0 | 0 | 0 | 3 | 6 | 29 | 17 | 50 | 11 | 6 | 0 | 0 | 0 | 0 | 0 |
| TMEM Tg B | 0 | 0 | 2 | 8 | 20 | 23 | 6 | 18 | 15 | 14 | 7 | 0 | 0 | 0 | 0 |
| d-TMEM Tg C | 0 | 0 | 0 | 0 | 1 | 3 | 5 | 35 | 34 | 24 | 5 | 0 | 0 | 0 | 0 |
| d-TMEM Tg D | 0 | 0 | 0 | 0 | 5 | 13 | 10 | 32 | 36 | 19 | 4 | 0 | 0 | 0 | 0 |
| dendra Tg | 0 | 0 | 0 | 0 | 0 | 0 | 0 | 0 | 1 | 2 | 17 | 66 | 10 | 2 | 0 |

| censors/day | 1 | 2 | 3 | 4 | 5 | 6 | 7 | 8 | 9 | 10 | 11 | 12 | 13 | 14 | 15 |
| --- | --- | --- | --- | --- | --- | --- | --- | --- | --- | --- | --- | --- | --- | --- | --- |
| N2 | 0 | 13 | 2 | 0 | 0 | 1 | 5 | 2 | 0 | 0 | 0 | 0 | 0 | 0 | 0 |
| TMEM Tg A | 0 | 0 | 0 | 0 | 0 | 0 | 0 | 0 | 0 | 0 | 0 | 0 | 0 | 0 | 0 |
| TMEM Tg B | 1 | 5 | 2 | 0 | 0 | 0 | 0 | 0 | 0 | 0 | 0 | 0 | 0 | 0 | 0 |
| d-TMEM Tg C | 0 | 1 | 1 | 1 | 1 | 0 | 0 | 1 | 0 | 0 | 0 | 0 | 0 | 0 | 0 |
| d-TMEM Tg D | 0 | 1 | 1 | 0 | 0 | 0 | 1 | 0 | 0 | 0 | 0 | 0 | 0 | 0 | 0 |
| dendra Tg | 0 | 0 | 0 | 0 | 0 | 0 | 0 | 0 | 0 | 1 | 0 | 0 | 0 | 0 | 0 |

Rep 3 (3/14-3/29)
| Deaths/day | 1 | 2 | 3 | 4 | 5 | 6 | 7 | 8 | 9 | 10 | 11 | 12 | 13 | 14 | 15 |
| --- | --- | --- | --- | --- | --- | --- | --- | --- | --- | --- | --- | --- | --- | --- | --- |
| N2 | 0 | 0 | 0 | 0 | 0 | 0 | 0 | 1 | 2 | 6 | 14 | 25 | 43 | 20 | 4 |
| TMEM Tg A | 0 | 0 | 0 | 5 | 18 | 24 | 18 | 23 | 14 | 9 | 10 | 1 | 0 | 0 | 0 |
| TMEM Tg B | 0 | 0 | 1 | 3 | 36 | 22 | 17 | 10 | 10 | 4 | 9 | 0 | 0 | 0 | 0 |
| d-TMEM Tg C | 0 | 0 | 1 | 9 | 10 | 16 | 11 | 23 | 16 | 7 | 7 | 0 | 0 | 0 | 0 |
| d-TMEM Tg D | 0 | 0 | 0 | 8 | 7 | 7 | 14 | 23 | 29 | 14 | 7 | 0 | 0 | 0 | 0 |
| dendra Tg | 0 | 0 | 0 | 0 | 0 | 0 | 1 | 2 | 2 | 3 | 24 | 51 | 23 | 8 | 0 |

| censors/day | 1 | 2 | 3 | 4 | 5 | 6 | 7 | 8 | 9 | 10 | 11 | 12 | 13 | 14 | 15 |
| --- | --- | --- | --- | --- | --- | --- | --- | --- | --- | --- | --- | --- | --- | --- | --- |
| N2 | 0 | 0 | 0 | 2 | 0 | 4 | 0 | 0 | 0 | 0 | 0 | 0 | 0 | 0 | 0 |
| TMEM Tg A | 3 | 0 | 0 | 0 | 0 | 0 | 0 | 0 | 0 | 0 | 0 | 0 | 0 | 0 | 0 |
| TMEM Tg B | 8 | 0 | 0 | 1 | 0 | 1 | 0 | 0 | 0 | 0 | 0 | 0 | 0 | 0 | 0 |
| d-TMEM Tg C | 2 | 0 | 7 | 1 | 1 | 0 | 0 | 0 | 0 | 0 | 0 | 0 | 0 | 0 | 0 |
| d-TMEM Tg D | 5 | 0 | 4 | 1 | 0 | 0 | 0 | 0 | 0 | 0 | 0 | 0 | 0 | 0 | 0 |
| dendra Tg | 0 | 0 | 2 | 0 | 0 | 0 | 0 | 2 | 0 | 0 | 0 | 1 | 0 | 0 | 0 |

## Notes

### Competing Interest Statement

The authors have declared no competing interest.

